# Novel murine models of post-implantation and midgestional malaria-induced preterm birth

**DOI:** 10.1101/2021.08.13.456287

**Authors:** Alicer K. Andrew, Caitlin A. Cooper, Julie M. Moore

**Affiliations:** Department of Infectious Diseases and Immunology, College of Veterinary Medicine, University of Florida, Gainesville, Florida, United States of America; Department of Cellular Biology, Franklin College of Arts and Sciences, University of Georiga, Athens, Georgia, United States of America; Department of Infectious Diseases, College of Veterinary Medicine, University of Georgia, Athens, Georgia, United States of America

## Abstract

Despite major advances made in malaria treatment and control over recent decades, the development of new models for studying disease pathogenesis remains a vital part of malaria research efforts. The study of malaria infection during pregnancy is particularly reliant on mouse models, as a means of circumventing many challenges and costs associated with pregnancy studies in endemic human populations. Here, we introduce three novel murine models that will further our understanding of how post-implantation and midgestional malaria infection affects pregnancy outcomes. When C57BL/6J (B6) mice are infected with *Plasmodium chabaudi chabaudi* AS on either embryonic day (E) 6.5, 8.5, or 10.5, preterm birth occurs in all animals by E16.5, E17.5, or E18.5 respectively, with no evidence of intrauterine growth restriction. We found that the time to delivery, placental inflammatory and antioxidant transcript upregulation, and the relationships between parasitemia and transcript expression prior to preterm birth are different based on the embryonic day of infection. On the day before preterm delivery, E6.5 infected mice did not experience significant upregulation of the inflammatory or antioxidant gene transcripts examined; however, parasitemia correlated positively with *Il1β*, *Cox1*, and *Hmox1* placental transcript abundance. E8.5 infected mice had elevated transcripts for *Ifnγ*, *Tnf*, *Il10*, *Cox1*, *Cox2*, *Sod1*, *Sod2*, *Cat*, and *Nrf2*, while *Sod3* was the only transcript that correlated with parasitemia. Finally, E10.5 infected mice had elevated transcripts for *Ifnγ* only, with a tendency for *Tnf* transcripts to correlate with parasitemia. Tumor necrosis factor deficient (TNF^-/-^) and TNF receptor 1 deficient (TNFR1^-/-^) mice infected on E8.5 experienced preterm birth at the same time as B6 controls. Further characterization of these models is necessary to discover new and/or existing mechanism(s) or trigger(s) responsible for malaria-driven preterm birth that is determined by gestational age upon maternal infection, parasite-host dynamics, or both.

## Introduction

Malaria remains a global public health threat despite unprecedented successes in vector control programs and antimalarial treatment. In 2019, the World Health Organization (WHO) reported that in regions with high to moderate *Plasmodium falciparum* transmission, roughly 11 million pregnancies were exposed to malaria infection [1]. Among those pregnancies, an estimated 872,000 infants were born with low birth weight, a well-documented risk factor for neonatal mortality [1–3]. Additional maternal-fetal health consequences to infection include maternal anemia and preterm delivery, especially in primi- and secundigravid women [4,5]. Notably, malaria infection during pregnancy can manifest as an organ-specific syndrome known as placental malaria (PM), identified by the accumulation of *Plasmodium*-infected red blood cells (IRBCs), immune cell infiltration, and both malaria pigment (hemozoin) and fibrin deposition in the placenta [6–9]. To protect pregnant women from PM, WHO recommends the use of insecticide-treated mosquito nets (ITNs) and intermittent preventative treatment using antimalarial drugs such as sulfadoxine-pyrimathamine (IPTp-SP). However, in 2018 only 31% of women received the recommended dosages of IPTp-SP during their pregnancy and ITN coverage in sub-Saharan Africa has stalled since 2015 [1]. Despite decades of discovery and billions of dollars invested in malaria research, the underlying mechanisms involved in PM pathogenesis are still incompletely understood. One major reason is that access to the placenta is limited until after delivery, making it difficult to evaluate the impact of infection on the placenta during earlier stages of pregnancy. As a result, most of what is known about human PM comes from studies focused on clinical outcomes throughout gestation, systemic evaluation for biomarkers of disease, and placental biopsies postpartum. Some studies provide evidence for the crucial role of inflammation, specifically via the upregulation of tumor necrosis factor (TNF), interleukin-10 (IL-10), interleukin-1 beta (IL-1β), and interferon-gamma (IFN-γ), in driving poor pregnancy outcomes [8,10–17]. Other studies suggest that oxidative stress, resulting from a cell’s inability to mount an effective antioxidant response to mitigate damage driven by reactive oxygen species (ROS), is a key player in the placental pathology and poor fetal health outcomes caused by PM [18,19].

Mouse models have been central to furthering our understanding of PM by providing a more accessible and genetically tractable tool for recapitulating PM development after infection in early pregnancy (pre-implantation) and late pregnancy (third trimester). For studies in early pregnancy, infection with the murine-infective *Plasmodium chabaudi chabaudi* AS (*Pcc*) in C57BL/6J (B6) and Swiss Webster mice capture some of the important hallmarks of human PM, such as elevated proinflammatory cytokine production, accumulation of IRBCs in the placenta, increased fibrin deposition, and pregnancy loss [20–25]. In these models, mice infected with *Pcc* on the first day of pregnancy (embryonic day 0.5, E0.5) experience a non-lethal infection that is accompanied by severe maternal anemia, high parasite burden, elevated inflammatory responses, hemozoin accumulation, and lipid peroxidation in the placenta, a known sign of oxidative stress. Therapeutic interventions that target the pathogenic contributions of coagulation, TNF, and ROS in B6 mice protect against pregnancy loss at midgestation (E10.5-12.5) [8,22,24]. Alternatively, studies in late pregnancy commonly utilize *Plasmodium berghei* in either B6 or BALB/c mice; however, the infection must be initiated between E10.5-13.5, due to the maternal lethality of the parasite. In these models, mice experience elevated placental inflammation and fibrin deposition, oxidative stress, IRBC adherence to the placenta, and preterm delivery [13,26–28].

Both antimalarial and anti-inflammatory drug treatment restore maternal survival, reduce oxidative stress, and improve pregnancy outcomes in these late pregnancy models [13,29]. While these well-established models are useful for studying PM pathogenesis, to our knowledge there have not been models reported that allow for comparative studies of non-lethal infection across distinct time points during mid-pregnancy. This presents an important obstacle in PM research because infection in the 2^nd^ trimester of human pregnancy, which is analogous to midgestation in mice, is associated with an increased risk of PM development [30,31]. Here, we describe three mouse models using *Pcc* in B6 mice to evaluate how the timing of malaria infection during post-implantation to midgestation (E6.5, E8.5, or E10.5) impacts pregnancy outcomes. Preliminary analyses of these models imply that different pathogenic mechanisms may be involved in the outcome of preterm delivery. Hence, these models will allow for interrogation of PM pathogenesis across distinct stages of pregnancy and at various stages during an acute, malaria infection.

## Materials and Methods

### Mice

C57BL/6J (B6), TNFα-knockout (B6;129S-Tnf^tm1Gkl^/J), TNFRp55-deficient (Tnfrsf1a^tm1Mak^/J), and A/J mice were purchased from the Jackson Laboratory (Bar Harbor, ME) and maintained by brother-sister mating for a maximum of ten generations in the University of Georgia Coverdell Vivarium, following guidelines and regulations set forth by the University of Georgia Animal Care and Use Committee. All animals were supplied food (PicoLab® Rodent Diet 20: 5030, St. Louis, MO) and water ad libitum. Mice were adjusted to a 14-hour light/10-hour dark cycle and housed in conditions of 65-75 °F and 40-60% humidity.

### Parasites and infection monitoring

The following reagent was obtained through BEI Resource Repository, NIAID, NIH: *Plasmodium chabaudi chabaudi*, Strain AS, MR4-741, contributed by David Walliker. Parasites were maintained as frozen stock in accordance with supplier guidelines and passaged in A/J mice for the purposes of infecting experimental B6 mice. Peripheral parasitemia was assessed by flow cytometry with a method adapted from work published by Jimenez-Diaz *et al.* [32]. A 2μl blood sample was collected by tail clip [33], diluted in 98μl 0.9% NaCl and stained with 0.25μl SYTO-16 Green Fluorescent Nucleic Acid Stain (ThermoFisher Scientific, catalog # S7578) within 4 hours of collection. Stained samples were incubated in the dark for 20 minutes at room temperature, further diluted 1:9 in 0.9% NaCl, then analyzed using a CyAn ADP Flow Cytometer (Beckman Coulter; Brea, CA). 30,000 cells were assessed daily for each mouse; infected red blood cells were distinguished based upon size and fluorescence intensity. An uninfected blood sample was used as an internal negative control. Parasitemia is reported as the percentage of infected red blood cells (IRBCs) to the total number of red blood cells (RBCs).

### Experimental Design

Female B6 mice aged 8-10 weeks were paired with age-matched males nightly and examined each morning until a vaginal plug was observed, indicating successful mating. The morning a vaginal plug was observed was considered embryonic day 0.5, E0.5. After baseline measurements of weight and hematocrit were recorded, females were left undisturbed until E 6.5 to minimize stress and increase the chances of successful blastocyst implantation. Mice were infected intravenously with 1000 *P. chabaudi chabaudi AS*-iRBCs diluted in 200ul 1X phosphate-buffered saline (PBS) per 20 grams of body weight on E6.5, E8.5, or E10.5 and are termed infected pregnant (IP). Age-matched non-pregnant mice were infected similarly as infection controls (infected non-pregnant, INP). In another control group, uninfected pregnant (UP) mice were sham injected with 200ul PBS per 20 grams body weight on the same gestation days. Immediately prior to infection or sham infection, experimental animals were switched to a high-fat rodent chow (PicoLab Mouse Diet 20 5058; St. Louis, MO) suited for pregnant animals. Weight measurements were recorded on E0.5, E6.5, E8.5, E10.5, and E12.5-E18.5 for all groups, depending on when the infection was initiated, to assess pregnancy progress and allow mice to proceed to spontaneous delivery. Parasitemia and hematocrit (a measure of anemia) were monitored daily in the infected groups beginning five days post-infection to assess the development of infection.

In a second series of experiments, mice infected on E6.5, E8.5, and E10.5, and along with their UP controls, were sacrificed on E15.5, E16.5, and E17.5, respectively. Weight, hematocrit, and parasitemia were recorded as described above until euthanasia. Mice were anesthetized with 2.5% Tribromoethanol before sacrifice and placentae were collected and preserved for histological and quantitative real-time PCR (RT-qPCR) analysis. Plump, pink, well-vascularized pups that reflexively responded to touch were considered viable and their weights and placenta weights were collected. Pup viability data are summarized in Supporting Table 1 (S1 Table).

### Histology

Placentae collected at the time of sacrifice were fixed for 24 hours in 10% buffered formalin, processed, and paraffin-embedded. Placental sections 5μm in thickness were mounted to microscope slides for histological analysis. Sections were Giemsa-stained and parasite burden in the placenta was determined by counting at least 1,000 erythrocytes in maternal blood sinusoids of at least two or more different placentae per dam as previously described [21].

### Gene expression by quantitative real-time PCR

Total RNA from mouse placentae collected on E15.5, E16.5, and E17.5 was isolated using Trizol Reagent (Ambion, Ref # 15596026) and a bead shaker (BeadBlaster 24, Benchmark Scientific, SKU: D2400) with a minimum of four placentae were pooled per dam. RNA was DNase-treated (Invitrogen, Ref # AM1906) and then reverse-transcribed with High-Capacity cDNA Reverse Transcription Kit (Applied Biosystems, Ref # 4368814). Relative transcript abundance for the genes of interest was quantified using PowerSYBR Green PCR Master Mix (Applied Biosystems, Cat # 4367659), the Roche LightCycler 96 Instrument (software version 1.01.01.0050), and the Mic qPCR cycler (Biomolecular systems, firmware version 2.25). Each sample was assayed in duplicate for target and housekeeping genes. Average Ct values of target genes were normalized to average Ct values of *Ubc* as the reference gene and relative transcript abundance of genes of interest was determined using the ∆∆Ct method. Transcript expression in individual mice is presented relative to the mean expression value in UP mice at E6.5. Details of primer sets are summarized in Supporting Table 2 (S2 Table).

### Statistics

All statistical analyses were performed using GraphPad Prism version 8.4.3 (GraphPad Software; La Jolla, California). All raw clinical data are presented as mean ± SEM. Error bars are not visible if they are shorter than the symbol’s height. The area under the curve (AUC) of percent starting weight, hematocrit, and parasitemia was calculated for each mouse between E0.5 and E18.5, as appropriate. AUC for weight and hematocrit was compared between IP, INP, and UP mice infected on the same day and was analyzed using a Kruskal-Wallis test between IP and INP groups and IP and UP groups. AUC for parasitemia between IP and INP groups was compared using a two-tailed unpaired t-test with Welch’s correction. RT-qPCR data were analyzed using an unpaired t-test with Welch’s correction and presented as a scatterplot with a bar representing the mean. Parasitemia and transcript data for correlation analyses were log-transformed. P values less than or equal to 0.05 were considered statistically significant. Mixed linear models analysis (SAS 9.4) was used to estimate differences in fetal and placental weights in dams sacrificed on E15.5, E16.5, and E17.5, and their controls. Each infection group was considered separately. The interrelatedness of placentae and pups born to the same dam was controlled with a random term for the dam. In each case, model fit with covariates considered to be relevant (infection status (binary variable), pup viability (as a proportion), and the total number of pups in each dam (continuous variable)) were compared against a model with no covariates. Proportional analysis tested by chi-square was used to compare pup viability between IP and UP dams.

## Results

### Infection at post-implantation and midgestation universally precipitates preterm pregnancy loss

In this observational study, *Pcc* blood-stage infection was initiated on E6.5, 8.5, or 10.5 in B6 mice. Initial measurements of weight and hematocrit were taken on E0.5 and then mice were randomly assigned to one of six groups – E6.5 IP, E8.5 IP, E10.5 IP, E6.5 UP, E8.5 UP, and E10.5 UP, with the indicated embryonic day representing the day of infection for IP groups (Fig 1a). After the infection was initiated, daily measurements of weight and parasitemia were collected beginning 5 days post-infection (Fig 1b-g). In each group, IP mice steadily gained weight alongside their UP controls until they experienced a sudden and precipitous decline in weight, indicative of preterm pregnancy loss (Fig 1b-d). The time to preterm birth was accelerated the later in gestation that infection occurred such that E6.5-, E8.5-, and E10.5- infected dams began losing weight nine, eight, and seven days post-infection respectively (Fig 1b-d). UP mice continued gaining weight beyond E18.5, as expected during a normal pregnancy. Parasitemia was greater in IP dams compared to their INP counterparts (Fig 1e-g), which was confirmed statistically through area under the curve (AUC) analysis, in all groups except the E8.5 infection group (Fig S1d-f). Corresponding with higher parasite burdens, AUC for hematocrit throughout the study was significantly lower in E6.5 infected dams compared to UP controls (Fig S1g), likely due to the prolonged destruction of red blood cells caused by the advanced stage of infection compared to other infection groups [34]. AUC for hematocrit in the other infection groups, E8.5 and E10.5, was not statistically significantly different from UP controls (Fig S1h-i).

**Fig 1.**
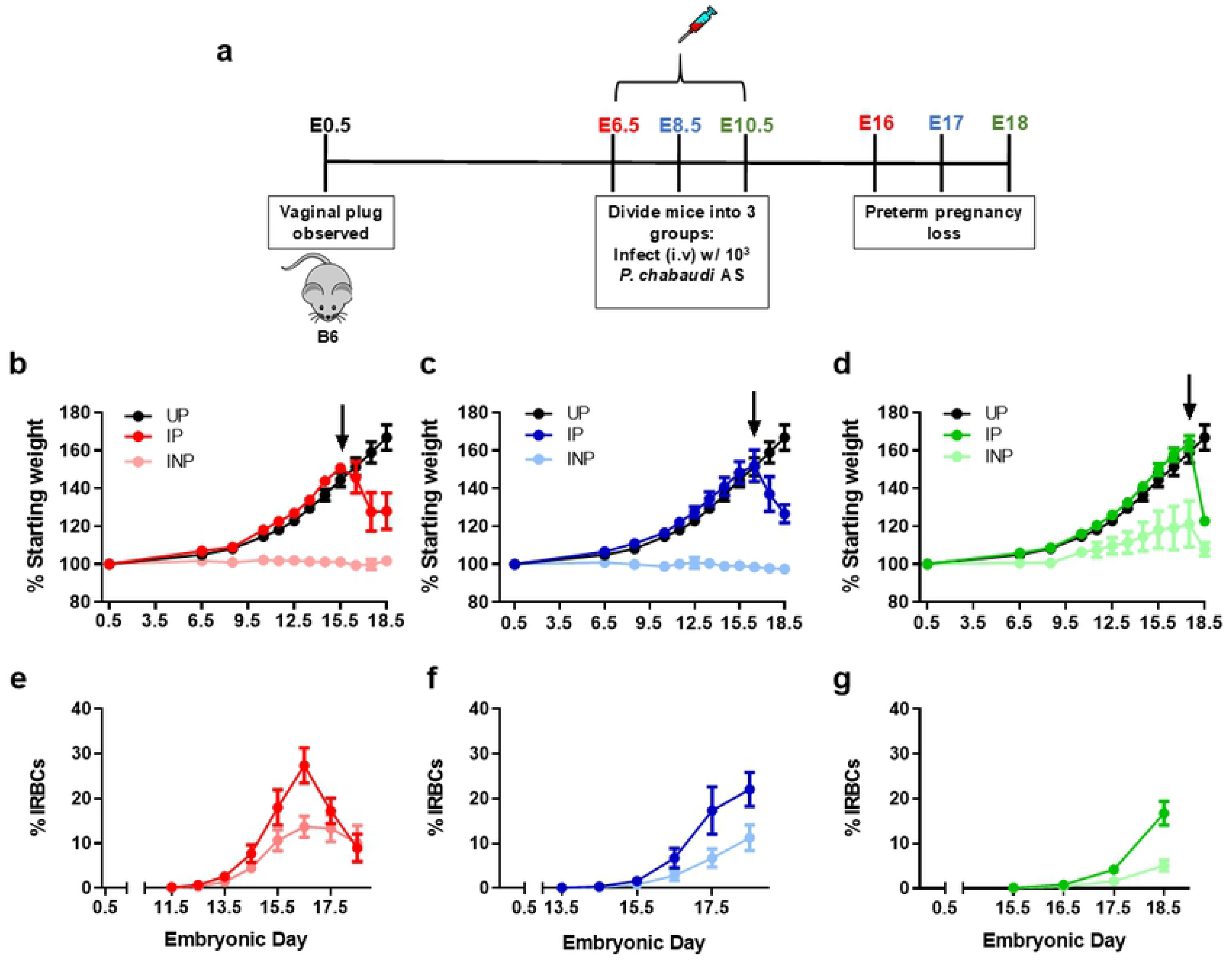
*P. chabaudi* AS infection from post-implantation to midgestation universally precipitates pre-term birth. Percent starting weight (a-c) and parasitemia (% IRBCs; d-f) are presented for infected pregnant (IP), uninfected pregnant (UP), and infected non-pregnant (INP) groups. All IP mice experienced precipitous weight loss, indicating pre-term delivery. Arrows indicate timepoints chosen for subsequent serial sacrifice experiments. (a, d) E6.5 infection group (red): IP, n=6; UP, n=6; and INP, n=4. (b, e) E8.5 infection group (blue): IP, n=8; UP, n=6; INP, n=4. (c, f) E10.5 infection group (green): IP, n=14; UP, n =6; INP, n=7.

### Placental parasitemia is elevated after infection but with no impact on placental or pup weight

To investigate the effect of maternal infection on placental and pup health in these models, a serial sacrifice study was conducted. Dams infected on E6.5, E8.5, or E10.5 were euthanized one day prior to expected preterm parturition on E15.5, E16.5, and E17.5 respectively (Fig 1b-d, arrows indicate the day of euthanasia when placentae were collected for analysis). Following infection, daily weight, parasitemia, and hematocrit measurements were recorded, beginning 5 days post-infection (Fig S2; Fig S3). Placental parasitemia in dams of the E8.5 and E10.5 infection groups, but not those infected at E6.5, was significantly higher than their peripheral parasitemia at sacrifice (Fig 2). To further examine the impact of infection, pup viability, placental weights, and pup weights were measured one day before expected preterm parturition (Fig 3; S1 Table). Pup viability, which was determined by the presence of a spontaneous reflex to touch, was not different between pups from IP versus UP dams, regardless of the infection group (Table S1). By mixed linear models analysis, neither placental nor pup weights were significantly impacted by maternal malaria across all infection groups; thus, graphs are shown for visualization purposes only (Fig 3). Holding infection status constant, increased pup viability in mice infected at E10.5 yielded a significant reduction in pup weight (by 0.7546 g, P = 0.0102), consistent with the well-documented observation that increased litter sizes naturally result in reduced fetal weights [35]. Otherwise, pup viability was unchanged between the pups from IP and UP dams in both the E6.5 and E8.5 groups. Finally, placental weights were not statistically different between IP and UP mice across all groups, however; placenta weights for both IP and UP mice in the E10.5 infection group were lower compared to the other groups, a phenomenon that occurs normally, as growth in placental weight and volume stagnate after E16.5 in mice [36,37]. Overall, these results indicate that despite having significant parasite burden, including in the placenta, malaria-induced preterm birth in these models occurs independently of changes in fetal and placental weight.

**Fig 2.**
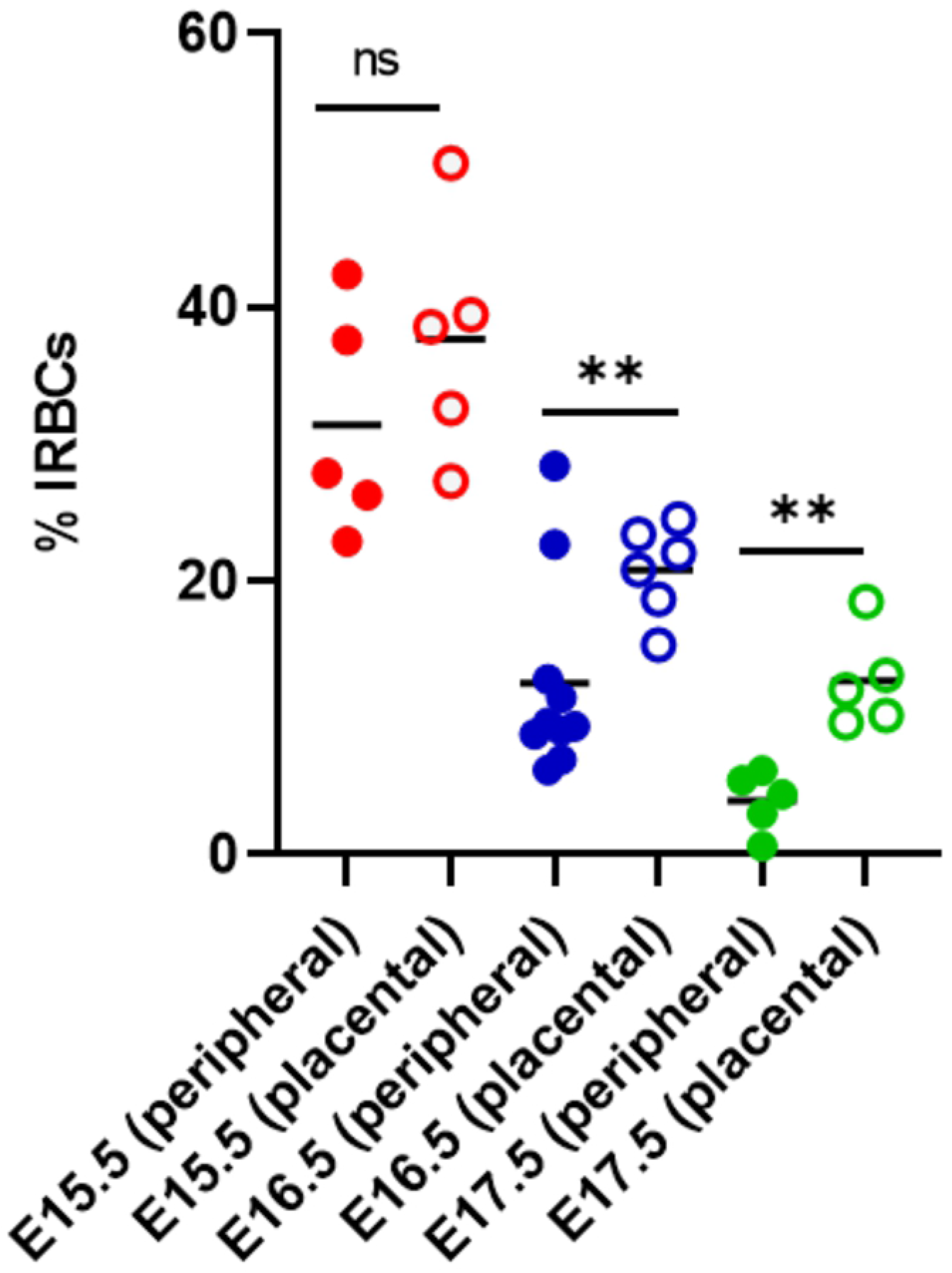
*P. chabaudi* AS-infected red blood cells accumulate in the placenta of mice in the E8.5 and E10.5 infection groups prior to preterm delivery. Peripheral and placental parasitemia from serially sacrificed mice are depicted. E15.5 placenta from E6.5 infected group (red), E16.5 placenta from E8.5 infection group (blue) and E17.6 placenta from E10.5 infection group (green). **P < 0.002, unpaired t-test with Welch’s correction; ns = not significant, P > 0.05.

**Fig 3.**
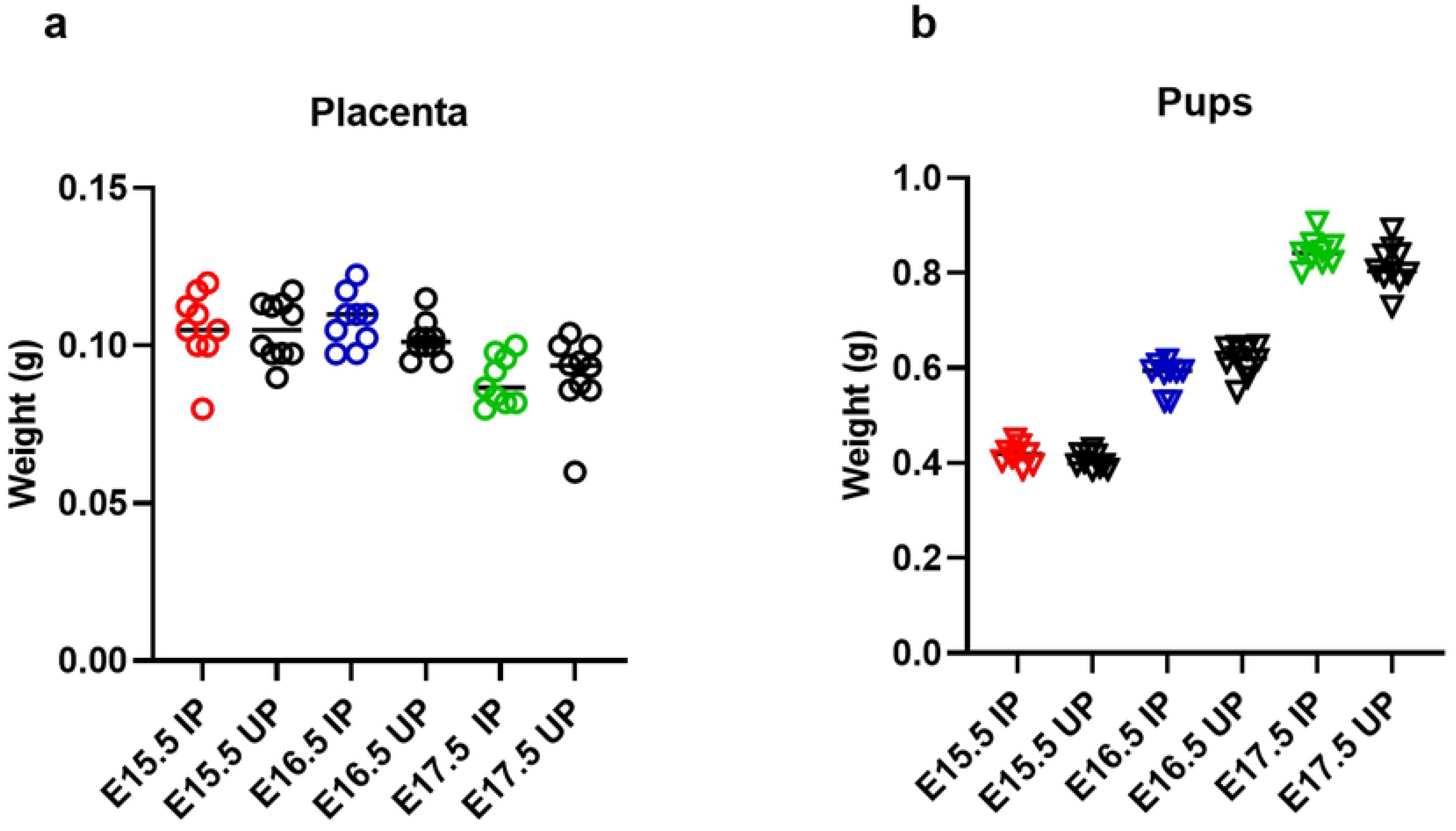
*P. chabaudi* AS infection does not significantly impact placenta or pup weight prior to pre-term delivery. Placenta (a) and pup weights (b) depicted for infected pregnant (IP) and uninfected pregnant (UP) mice sacrificed one day before preterm delivery. These data are shown for visualization only; statistical analysis is in the text. E6.5 infection group: 4 IP dams, 32 placentae; 4 UP dams, 30 placentae. E8.5 infection group: 4 IP dams, 31 placentae; 4 UP dams, 33 placentae. E10.5 infection group: 5 IP dams, 40 placentae; 5 UP dams, 41 placentae.

### Placental inflammatory responses do not universally precede preterm delivery

The pathogenesis of preterm delivery has long been associated with excessive inflammation, induced by infection or other disorders [38]. To probe the importance of inflammation in driving preterm delivery in these models, real-time quantitative PCR (RT-qPCR) analysis was performed using RNA isolated from the placental homogenates from serially sacrificed dams. The relative abundance of inflammatory gene transcripts, such as tumor necrosis factor (*Tnf*), interleukin-10 (*Il10*), interferon-gamma (*Ifnγ*), interleukin beta (*Il1β*), and genes important in pregnancy maintenance and parturition, such as cyclooxygenase 1 and 2 (*Cox1* and *Cox2*) were measured (Fig 4). Transcript abundance in placentae collected on E15.5 from mice infected on E6.5 revealed no significant differences between IP and UP dams for all inflammatory targets measured (Fig 4a-f). However, *Il1β* transcripts were positively correlated with both peripheral parasitemia at the time of sacrifice (E15.5) and parasitemia AUC for mice infected on E6.5 (Fig 5a-b). Additionally, there was a positive correlation between *Cox1* transcripts and parasitemia at sacrifice (Fig 5c) and a weak tendency for a positive correlation between *Cox1* transcripts and parasitemia AUC for this same group (Fig S4a). In the E8.5 infection group, transcript abundance of all the inflammatory genes targeted, except for *Il1b*, was significantly elevated in placentae collected on E16.5 (Fig 4a-f), while no correlative relationships with parasitemia at sacrifice or parasitemia AUC were found. In the E10.5 infection group, only transcripts for *Ifng* were elevated, while all other targets remained unchanged (Fig 4a-f). However, there was a tendency for *Tnf* transcripts to be positively correlated with both peripheral parasitemia at sacrifice and parasitemia AUC in mice infected on E10.5 (Fig S4b-c).

**Fig 4.**
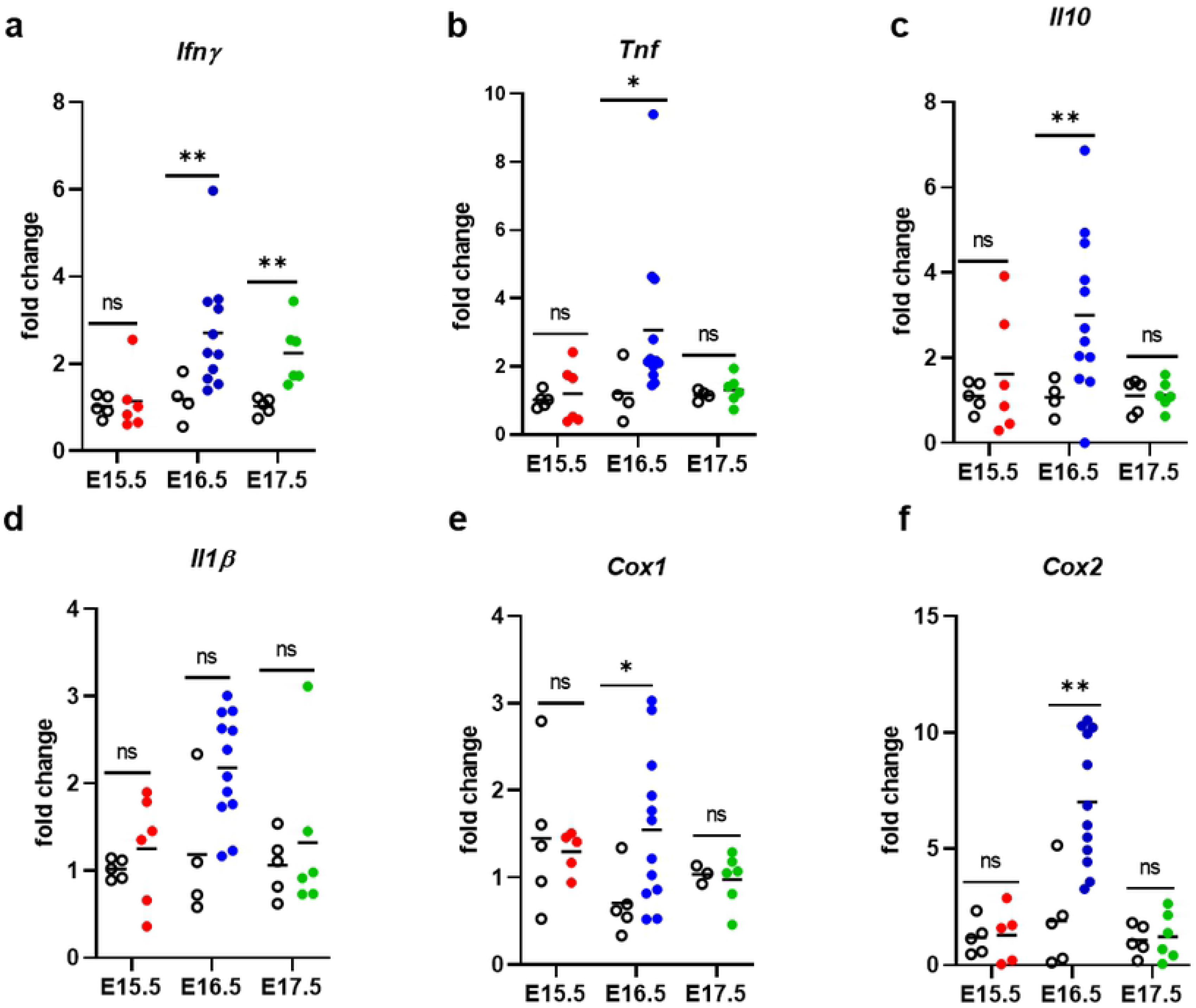
*P. chabaudi* infection is associated with inflammatory and parturition-associated gene transcript upregulation in a gestational day-dependent manner one day before preterm delivery. Mouse *Ifnγ*, *Tnf*, *Il10*, *Il1β*, *Cox1,* and *Cox2* mRNA abundances (a-f) were normalized to *Ubc* and quantified by qPCR in infected pregnant (solid circles) and uninfected pregnant (open black circles) placentae. Colored solid circles denote E15.5 placenta from the E6.5 infected group (red), E16.5 placenta from the E8.5 infection group (blue), and E17.6 placenta from the E10.5 infection group (green). Group means and transcript abundance in individual mice are depicted. **P ≤ 0.005, *P < 0.05, unpaired t-test with Welch’s correction; ns = not significant, P > 0.05.

**Fig 5.**
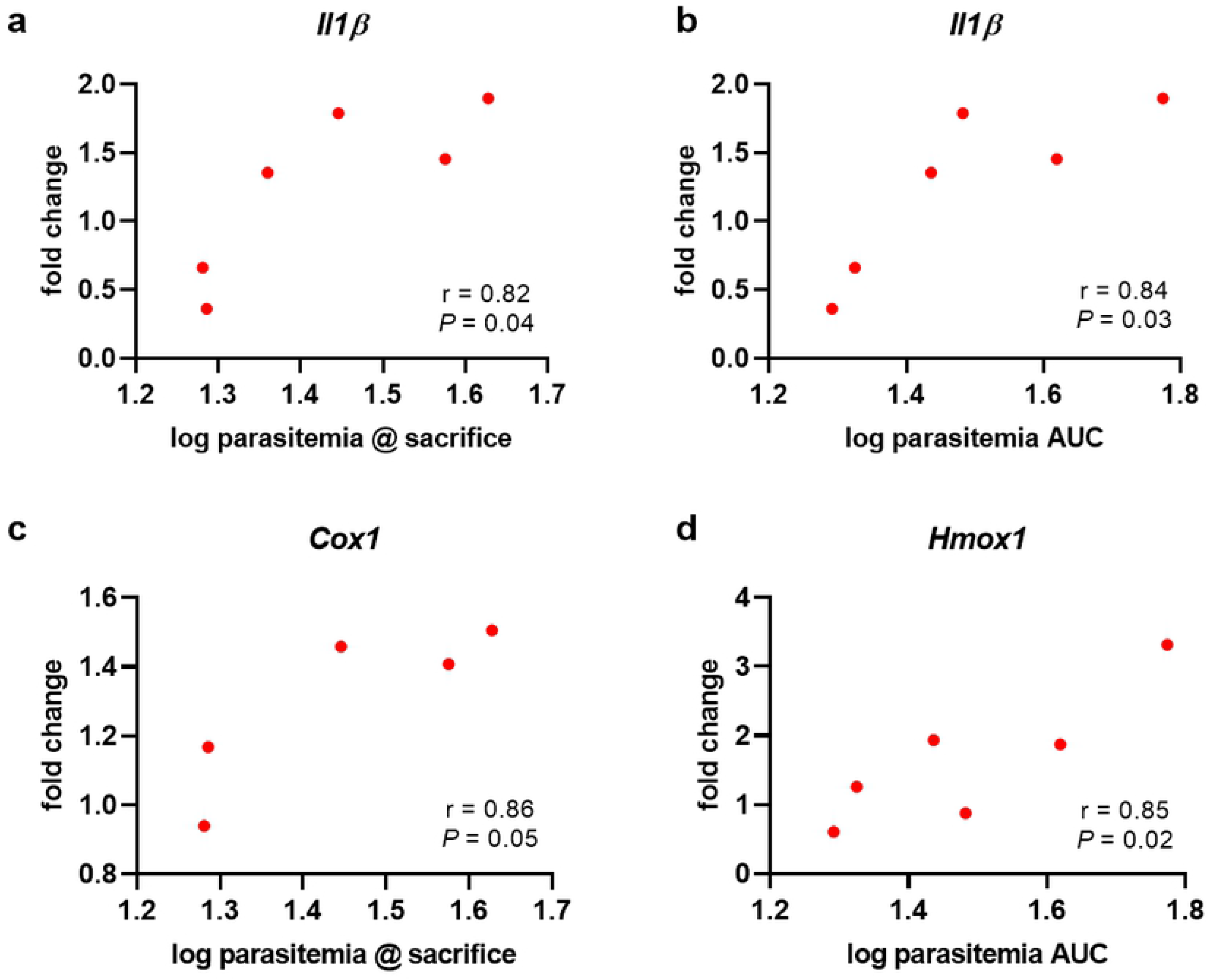
Inflammatory and antioxidant gene expression positively correlates with parasitemia in mice infected on E6.5. Mouse mRNA transcript abundance relative to parasitemia at the time of sacrifice or parasitemia AUC in placenta collected one day before expected pre-term birth (E15.5). *IL1β* transcripts were positively correlated with both parasitemia at the time of sacrifice (a) and parasitemia AUC (b). *Cox1* transcripts were positively correlated with parasitemia at the time of sacrifice (c) and *Hmox1* transcripts were positively correlated with parasitemia AUC.

To directly evaluate a link between placental inflammation and precipitation of preterm delivery at E16.5, mice deficient in tumor necrosis factor (TNF^-/-^) and tumor necrosis factor receptor 1 (TNFRI^-/-^) were infected with 1000 *Pcc*-IRBCs on E8.5. Both TNF^-/-^ and TNFRI^-/-^ IP mice experienced preterm delivery beginning on E17.5, similar to wildtype B6 mice (Fig 6). Although parasitemia appears to be lower in TNF^-/-^ IP compared to B6, AUC analysis revealed no significant differences between the groups for either TNF^-/-^ or TNFRI^-/-^ IP dams (Fig S5). Taken together, these data demonstrate that TNF and TNF signaling through TNFRI are not required to drive preterm delivery in the E8.5 infection model.

**Fig 6.**
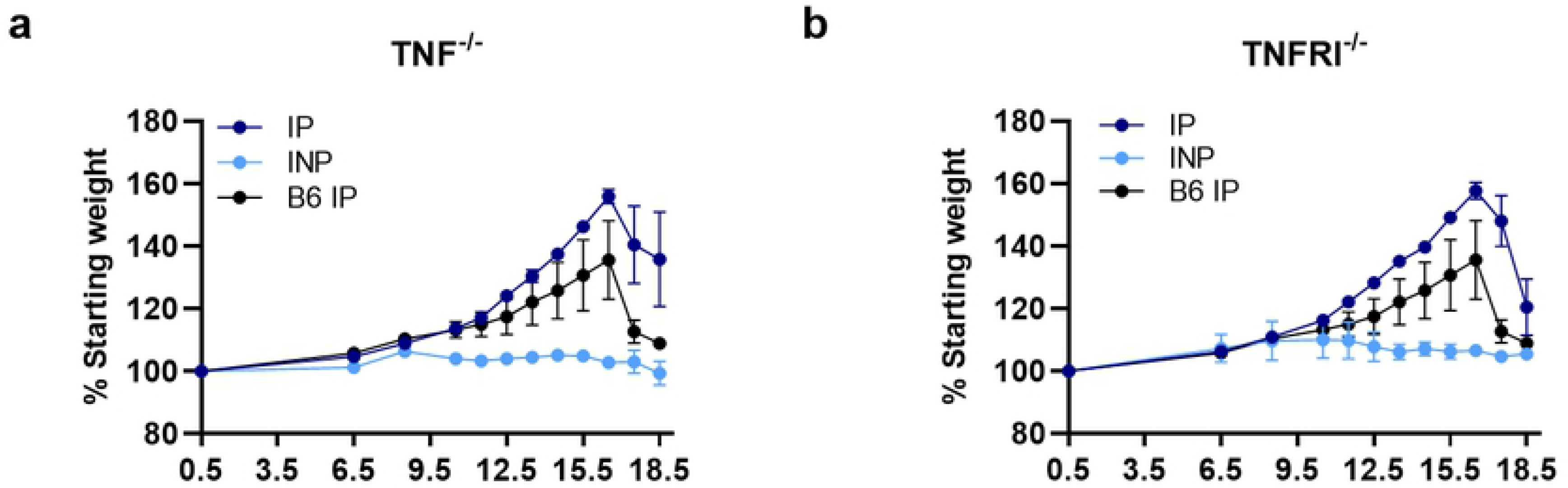
Tumor necrosis factor (TNF^-/-^) and tumor necrosis factor receptor 1 (TNFRI^-/-^) deficient mice experience pre-term birth after *P. chabaudi* infection on E8.5. Percent starting weight for TNF^-/-^ (a) and TNFRI^-/-^ (b) mice are depicted for infected pregnant (IP) and infected non-pregnant (INP) groups. TNF^-/-^ IP: n = 3, INP n =3; TNFRI^-/-^ IP: n = 7, INP n = 3; B6 IP: n = 5. All IP mice experience precipitous weight loss, indicating pre-term delivery.

### Placental antioxidant responses do not universally precede preterm delivery

Oxidative stress, generally described as an imbalance between the formation of free radicals, such as reactive oxygen species (ROS), and protective antioxidant defense molecules within cells and tissues, leads to cellular damage to lipids, proteins, RNA, and DNA. Recently, oxidative stress has been documented to play a crucial role in pregnancy outcome in some mouse models of PM [24,26,27] and is implicated in human populations experiencing malaria during pregnancy [18,19] as well as other types of pregnancy complications [38–40]. To interrogate the role of oxidative stress in our models, placentae from serially sacrificed mice were evaluated for alterations in antioxidant gene expression by RT-qPCR (Fig. 7). In the placentae of B6 mice infected on E8.5, transcript abundance for superoxide dismutase 1 and 2 (*Sod1, Sod2*), catalase (*Cat*), and nuclear factor erythroid-2 related factor 2 (*Nrf2*) were significantly elevated compared to UP controls (Fig 7a-b, d-e). None of these targets were significantly elevated in IP mice infected on E6.5 or E10.5 (Fig 7a-f). However, when the relationship between antioxidant transcript abundance and parasitemia was considered, *Hmox1* was positively correlated with parasitemia AUC for mice in the E6.5 infection group (Fig 5d). A weak tendency for a positive correlation was found between *Cat* transcripts and parasitemia AUC and *Hmox1* transcripts and parasitemia at sacrifice (Fig S4 b-c). Additionally, *Sod3* transcripts were positively correlated with both parasitemia at sacrifice (Fig 8) and parasitemia AUC (r = -0.56, P = 0.05, n = 12; data not shown) in mice infected on E8.5. Significant correlations between antioxidant transcript expression and parasite burden were not found in the E10.5 infection group.

**Fig 7.**
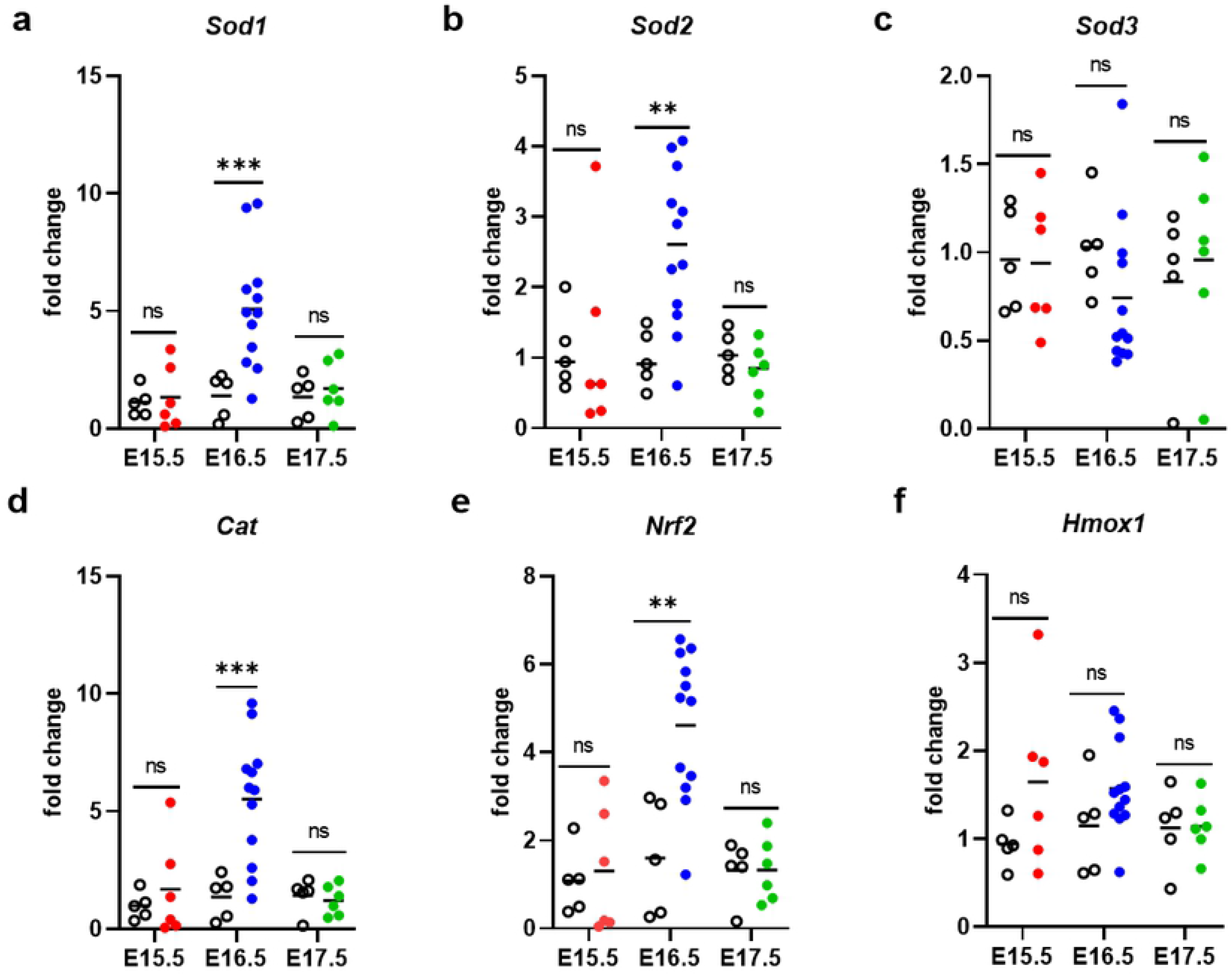
*P. chabaudi* infection on E8.5 leads to elevated mRNA transcript abundance for some antioxidant genes one day before preterm delivery. Mouse *Sod1*, *Sod2*, *Sod3*, *Cat, Nrf2, and Hmox1* mRNA abundance (a-f) were normalized to *Ubc* and quantified by qPCR in infected pregnant (solid circles) and uninfected pregnant (open black circles) placentae. Colored solid circles denote E15.5 placenta from the E6.5 infected group (red), E16.5 placenta from the E8.5 infection group (blue), and E17.6 placenta from the E10.5 infection group (green). Group means and transcript abundance in individual mice are depicted. ***P ≤ 0.005, **P = 0.02; ns = not significant, P > 0.05.

**Fig 8.**
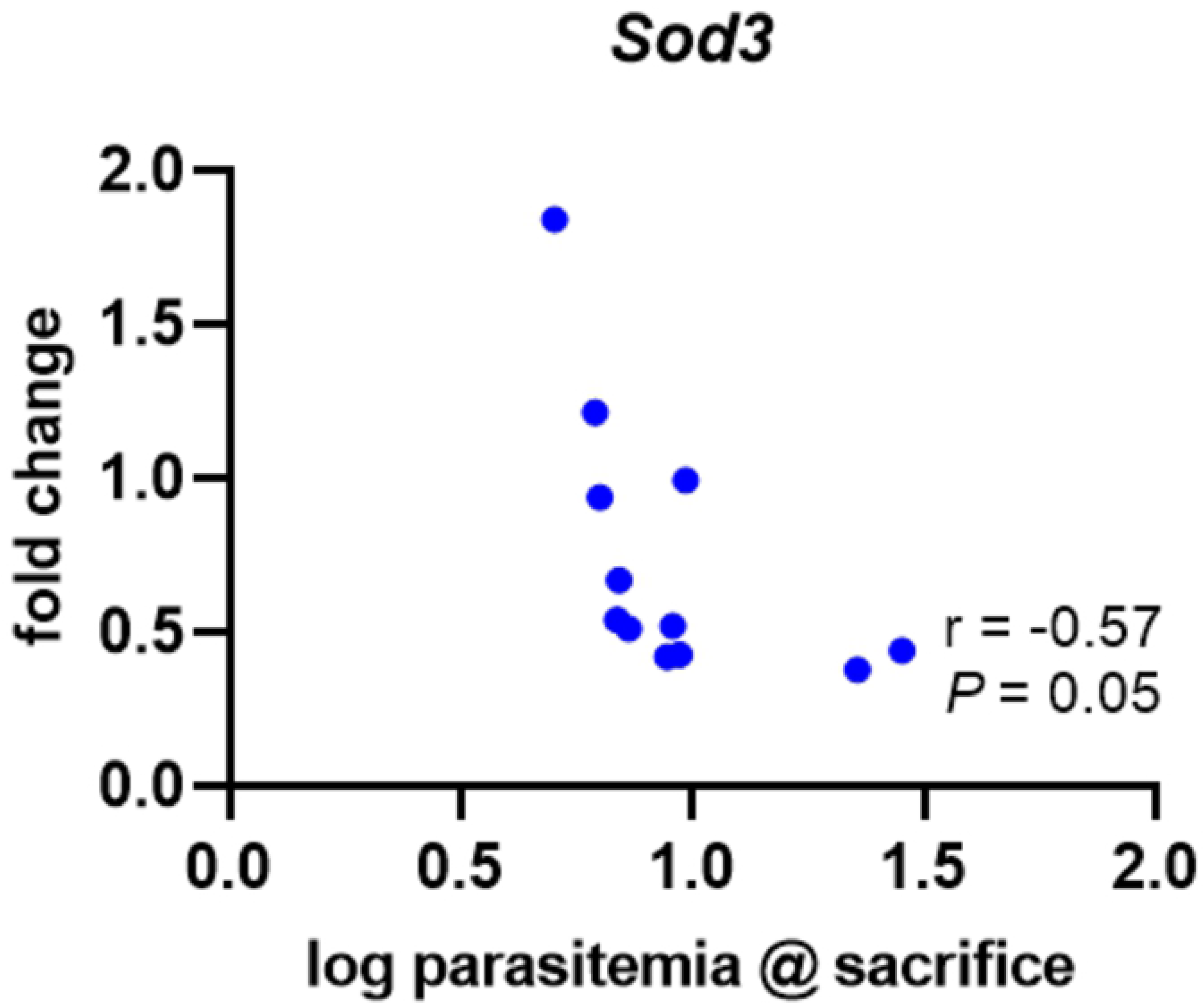
Antioxidant gene expression negatively correlates with parasitemia in mice infected on E8.5. Mouse *Sod3* mRNA transcript abundance relative to parasitemia at the time of sacrifice and measured in placentae collected one day before expected preterm birth (E16.5).

## Discussion

Mouse models represent an affordable and genetically manipulable tool for investigating the pathogenesis of PM, a syndrome responsible for significant maternal morbidity and poor pregnancy outcomes worldwide. In this study, three murine models for malaria-induced preterm birth were characterized. These models require a blood-stage infection in C57BL/6J (B6) mice using the non-lethal murine infective parasite, *Plasmodium chabaudi chabaudi* AS (*Pcc*) at E6.5, E8.5, and E10.5. Although post-implantation *Pcc* infection in these models universally results in preterm labor, the results show that transcript abundance for certain genes in the placenta prior to labor differ based on the model. To our knowledge, these three models, combined with initiation of infection at E0.5 [21,22,25] are the first to provide the flexibility to probe the effect of malaria infection spanning the pre-implantation to the midgestation period in the mouse, without inducing maternal mortality. To varying extents, key features of human PM are recapitulated in these models, including severe maternal anemia, higher parasitemia in the placenta compared to the periphery, and preterm delivery [8,20–22,24,25]. Indeed, in our 8.5 and 10.5 infection models, higher parasitemia in pregnant mice compared to non-pregnant controls corresponds with higher parasitemia in the placenta rather than the periphery, corroborating previous studies in both mice and humans demonstrating that pregnancy itself can increase susceptibility to malaria [4,27,28,41] and lead to parasite accumulation in the placenta [9]. Thus, these models provide an opportunity for researchers to take a snapshot of the placenta at various times during the course of an acute malaria infection to investigate the relationships between parasite dynamics, gestational age, and birth outcome.

Preterm delivery was a universal outcome of infection in these new models, with the time to delivery shortening the later in gestation the infection was introduced. When placenta weight, pup weight, and pup viability were assessed one day prior to preterm delivery, the infection status of the dam had no impact. Thus, the physiological events that precipitate labor are acute. Analysis of the placenta preceding preterm delivery revealed differential expression of various inflammatory gene transcripts, highlighting the possibility that distinct mechanisms or triggers may be involved in driving preterm delivery in a gestational age-dependent manner. Studies of cyclooxygenases, COX1 and COX2, have described their importance in the onset of labor in both mice and humans [42,43]; however, *Cox1* and *Cox2* transcripts were elevated in the E8.5 infection model only. *Cox1* was positively correlated with parasitemia in the E6.5 infection model but no significant malaria-induced upregulation for either *Cox1* or *Cox2* was detected in the E10.5 infection model one day prior to preterm delivery. Moreover, when *Cox1* and *Cox2* expression was compared amongst the UPs from all experimental groups (data not shown); there was no relationship between transcript abundance and gestational age. These findings imply that the changes in *Cox1* and *Cox2* transcripts observed are both malaria-induced and dependent on the embryonic day of infection. IFNγ is elevated during acute malaria infection and required for parasite control [44]. *Ifnγ* transcripts were only elevated in the E8.5 and E10.5 groups even though mice in all infection groups were experiencing ascending parasitemia at the time that placental analyses were conducted. Additionally, *Tnf* and *Il10* transcripts were increased in the E8.5 infection group only, despite being associated with human malaria-induced preterm delivery [10,17]. Interestingly, TNF^-/-^ and TNFR^-/-^ mice infected on E8.5 did not experience improved pregnancy outcomes or any changes in the time to preterm delivery. These data suggest that although TNF signaling may be an important driver of poor pregnancy outcomes during malaria infection [10,11,22], it is not directly involved in driving preterm delivery in the E8.5 model. Additional experiments in which TNF^-/-^ and TNFR1^-/-^ mice are infected on E6.5 and E10.5 are required to fully determine the role of TNF in both infection control and pregnancy outcomes in those models. *Il1β* transcript abundance positively correlated with parasitemia in the E6.5 infection group only but no changes in its expression were detected in the other infection models. This observation is noteworthy because in another model of infection during late pregnancy, reduction of IL-1β signaling improves pregnancy outcome [13]; thus, IL-1β may play a pathogenic role in our E6.5 infection model as well, perhaps through inflammasome assembly and/or the initiation of pyroptosis in gestational tissues, as has been demonstrated in other models of PM [45].

Since oxidative stress has been described in various models of preterm delivery [46] and implicated in both human and mouse PM [18,19,24,26,27], antioxidant gene transcripts were measured in the placenta one day before preterm delivery. *Sod1*, *Sod2*, *Cat*, and *Nrf2* transcripts were elevated in the E8.5 infection group only. These data are further corroborated by correlation analyses which show that *Sod3* transcripts are positively correlated with both parasitemia at sacrifice and parasitemia AUC in this same infection group. Similar results were reported in another model of PM, where antioxidant gene transcripts were elevated in B6 conceptuses and antioxidant drug treatment helped to mitigate pregnancy loss [24]. Since none of the targeted antioxidant genes were elevated in the placenta of the E6.5 and E10.5 infection groups prior to preterm delivery, it may imply that other biological processes, not oxidative stress, will be determined as more suitable therapeutic targets for pregnancy loss when an infection is initiated at these timepoints specifically. On the other hand, *Hmox1* transcripts were positively correlated with parasitemia AUC in the E6.5 infection group, but not in the other models. Since E6.5 IP mice exhibited the longest time to preterm delivery compared to the other two groups, this particular model could present another opportunity to study the impact of *Hmox1* regulation of heme in driving negative pregnancy outcomes. Elevated heme levels and altered *Hmox1* activity have already been implicated in the pathogenesis of both human and murine PM [47–49] and other pregnancy complications [50]; however, more research is required to identify any therapeutic potential in our model. For the E10.5 model, none of the antioxidant gene transcripts were significantly differentially expressed or correlated with parasite burden. Because this group overall had a lower parasite burden relative to the other groups, these results may collectively indicate that a threshold of infectious burden is required to galvanize oxidative damage and commensurate antioxidant responses.

Although the characteristics of these models are largely descriptive thus far, they may be instrumental for future studies aimed at examining maternal and/or fetal responses to malaria infection at various times during gestation. Interestingly, while infection at different stages of pregnancy ultimately ends in the same outcome, preterm birth, distinct pathological pathways to achieving this result may be involved. Indeed, there are reports of differential expression of inflammatory mediators such as IL-10 and toll-like receptors and altered gene methylation patterns in the human placenta across various times during gestation [51–53]. Additionally, proper murine placental function can be sensitive to the timing of insult during gestation [54]. Given this knowledge, it is possible that in our models the placenta is demonstrating the ability to respond to malaria infection in a gestational age-specific manner that eventually ends in preterm delivery. On the other hand, the timing of our analyses may have led us to miss critical physiological changes that occurred closer to the time of labor. The presence of a universal trigger(s) that favors preterm delivery independently of any differences in placental maturity or development at the time of infection is also conceivable. Our analyses focused on placental inflammation and oxidative stress; however, there are many other mechanisms of infection-induced preterm delivery that could be assessed in these models. Some biological functions of clinical relevance in PM that could be explored in future studies in these models include the coagulation pathway [8,55], the complement system [56], nutrient bioavailability [57] and transport across the placenta [58,59], angiogenic balance and vascularization [60], placental autophagy [61,62], and hormonal alterations [61]. Finally, it is possible that pathological changes leading to preterm birth are occurring in tissue types that we did not evaluate, such as the plasma or uterus. Interactions at the maternal-fetal interface are associated with poor pregnancy outcomes in malaria [49,63]; thus, uteroplacental physiological changes that were not measured here may be critical. Moreover, the ability to successfully detect and characterize physiologically relevant changes by RT-qPCR alone may be limited. Thus, follow-up studies must consider expanding the types of tissues evaluated and the breadth of techniques used to identify a more precise cause(s) of preterm birth in these models.

In conclusion, the new models described here provide a new avenue for interrogating the pathological driver(s) of preterm birth and/or gestation-age-dependent malaria-induced pregnancy compromise following post-implantation and midgestation infection. These models expand our available tools for studying the mechanisms involved in PM pathogenesis and as such could help improve our understanding of how malaria during pregnancy may galvanize preterm delivery.

## Acknowledgments

We thank MR4 for providing the malaria parasites contributed by David Walliker. We thank Dr. Demba Sarr and Dr. Catherine D. Morffy Smith for providing the primers used for the RT-qPCR assays. Alicer K. Andrew was supported by the 2017-2019 Peach State LSAMP Bridge to the Doctorate Program at the University of Georgia (National Science Foundation, Award # 1702361). This work was supported by National Institute of Health grants to J.M.M. The content is solely the responsibility of the authors and does not necessarily represent official views of the *Eunice Kennedy Shriver* National Institute of Child Health and Human Development, the National Institute of Allergy and Infectious Diseases, or the National Institutes of Health.

## Supporting Information Captions

**Fig S1. Area under the curve analysis for weight, parasitemia, and hematocrit in B6 mice infected with *P. chabaudi* AS from post-implantation to midgestation.**

Area under the curve (AUC) was calculated for uninfected pregnant (UP), infected pregnant (IP) and infected non-pregnant (INP) mice belonging to the E6.5 infection group (red), E8.5 infection group (blue), or E10.5 infection group (green) in an observational study. No statistical differences are observed in weight change over time between UP and IP animals (a-c). Parasitemia AUC is higher in some IP animals compared to INP counterparts (d-f). Hematocrit AUC was statistically different in the E6.5 infection group only (g-i). Groups were compared either by using a Kruskal-Wallis test or unpaired t-test with Welch’s correction (for parasitemia AUC). **P ≤ 0.005, *P < 0.05; ns = not significant, P > 0.05.

**Fig S2. Course of *P. chabaudi* AS infection in mice sacrificed one day prior to expected pre-term delivery.**

Percent starting weight (a-c) and parasitemia (% IRBCs; d-f) are presented for infected pregnant (IP), uninfected pregnant (UP), and infected non-pregnant (INP) groups. All mice were serially sacrificed one day prior to expected preterm delivery and tissues were collected for further studies. (a, d, g) E6.5 infection group, euthanized on E15.5 (red): IP, n=9; UP, n=9; and INP, n=4. (b, e, h) E8.5 infection group, euthanized on E16.5 (blue): IP, n=15; UP, n=12; INP, n=7. (c, f, i) E10.5 infection group, euthanized on E17.5 (green): IP, n=12; UP, n =10; INP, n=6.

**Fig S3. Area under the curve analysis for serially sacrificed mice.**

Area under the curve (AUC) was calculated for uninfected pregnant (UP), infected pregnant (IP) and infected non-pregnant (INP) mice belonging to the E6.5 infection group (red), E8.5 infection group (blue), or E10.5 infection group (green) for serial sacrifice experiments. No statistical differences are observed in weight change over time between UP and IP animals (a-c). Parasitemia AUC is higher in some IP animals compared to INP counterparts (d-f). Hematocrit AUC achieved statistical significance between IP and INP mice in the E8.5 infection group only (g-i). Groups were compared either by using a Kruskal-Wallis test or unpaired t-test with Welch’s correction (for parasitemia AUC). ****P < 0.0001, **P < 0.005, *P < 0.05; ns = not significant, P > 0.05.

**Fig S4. Correlation analyses between transcript target and parasitemia in placentae of mice in the E6.5 and E10.5 infection groups.**

Mouse mRNA transcript abundance relative to parasitemia at the time of sacrifice or parasitemia AUC in placenta collected one day before expected pre-term birth. *Cox1* (a) and *Cat* (e) transcripts tended to be positively correlated with parasitemia AUC. *Tnf* (c) and *Hmox1* (d) transcripts displayed a similar tendency with parasitemia at sacrifice. *Tnf* transcripts (b, c) tended to be positively correlated with both parasitemia AUC and parasitemia at the time of sacrifice.

**Fig S5. Parasitemia and area under the curve for parasitemia in TNF^-/-^ and TNFR1^-/-^ mice infected on E8.5.**

Parasitemia (% IRBCs) in TNF^-/-^ (a) and TNFRI^-/-^ (b) mice. Area under the curve (AUC) analysis (c, d) do not show a statistically significant increase in parasitemia between IP versus INP groups for both strains. TNF^-/-^ IP: n = 3, INP n = 3; TNFRI^-/-^ IP: n = 7, INP n = 3; ns = not significant, P > 0.05.

**S1 Table. Pup viability between infected and uninfected dams sacrificed one day prior to expected preterm delivery.**

No differences were observed in pup viability between infected pregnant (IP) and uninfected pregnant (UP) dams across all infection groups. Statistical significance determined via proportional analysis tested by chi-square.

**S2 Table. Primer sequences for qPCR targets.**

Mouse-specific forward (FP) and reverse (RP) primers used in quantitative real-time PCR for the amplification of mRNA transcripts associated with inflammation, parturition, antioxidant activity, and reference (*Ubc*) genes.

